# *Six3* acts independently of *Pax6* to provide an essential contribution to lens development

**DOI:** 10.1101/2023.01.09.523289

**Authors:** Sumanth Manohar, Takuya Nakayama, Marilyn Fisher, Robert M. Grainger

**Affiliations:** Department of Biology, University of Virginia, Charlottesville, VA 22904, USA

**Keywords:** Six3, Pax6, lens induction, *Xenopus*, BMP signaling, presumptive retina

## Abstract

The Six3 transcription factor is essential for forebrain and eye development, and *SIX3* mutations cause the congenital disorder holoprosencephaly. We created a *six3* mutant in *Xenopus tropicalis* with a mild holoprosencephaly phenotype, and unlike mouse *Six3* mutants that are headless/eyeless, the *Xenopus* mutant forms some eye structures, allowing direct study of Six3 function in eye formation. We focus here on striking deficits in lens formation. Early lens induction occurs normally in the mutant, e.g., the essential eye gene *pax6*, is activated in lens ectoderm, persisting in the eye to a late developmental stage, but in many embryos the lens fails to form. We found that *bmp4, bmp7*.*1, smad7, dll1, dlc, mab21l1* and/or *mab21l2*, previously unknown as *six3* eye targets, are downregulated in the mutant. We show that *six3* is required for lens formation, acting primarily in developing retina during neurulation through BMP and Notch signaling, and that *mab21l1/mab21l2* regulate(s) this BMP activity. This work reveals previously unrecognized essential roles for *six3* in eye development, identifying its key role in signaling needed for lens formation, and acting independently of *pax6* activity.

**SUMMARY STATEMENT:** This study identifies the *six3* transcription factor as the mediator of key inductive signals driving lens formation, acting independently of *pax6* in early phases of lens formation.

## INTRODUCTION

Vertebrate eye formation is strongly linked to the gene *Pax6* (Gehring and Ikeo, 1999); however, studies have demonstrated contributions from the Six (sine oculis homeobox) family of transcription factors (TFs) to eye formation, both from studies in *Drosophila* and vertebrates (Kumar, 2009). The Six protein family possesses two characteristic domains, a Six domain (SD) required for protein-protein interaction and a DNA-binding homeodomain (HD). In *Drosophila* the *sine oculis* (*so*) gene is a *Six* gene family member that plays a key role in retinal gene circuitry, whereas in vertebrates the *Six3*/*Six6* subfamily of *Six* genes is involved in development of the eye and/or forebrain (Kawakami et al., 2000; Kumar, 2009).

In humans, *SIX3* mutations cause holoprosencephaly (HPE), a failure to properly form and separate left and right sides of the forebrain during embryonic development (Dubourg et al., 2007). We previously reported that targeted disruption of the *six3* gene via CRISPR/Cas9 gene editing resulted in HPE-like phenotypes in F0 *Xenopus tropicalis* embryos (Nakayama et al., 2013). The germline-transmitted *six3* presumptive null mutant of *X. tropicalis* (see below) also has an HPE phenotype and still undergoes early stages of eye formation. *Six3* null mice have a stronger phenotype than in human patients or *Xenopus*, resulting in absence of the rostral forebrain, the eyes, and nose leading to postnatal lethality (Lagutin et al., 2003), thus only permitting study of eye development using conditional knockout mutations targeting eye tissues (e.g., Liu et al., 2006, Liu et al., 2010). The latter generate mutant phenotypes generally induced after the earliest stages of eye formation, and lens-specific conditional targeting vectors also may affect lens gene regulation (Dorà et al., 2014; Pathania et al., 2016; Song and Palmiter, 2018).

*Six6* complements *Six3* activity (Diacou et al., 2018; Liu and Cvekl, 2017), and *SIX6* mutations cause pituitary anomalies and eye defects including anophthalmia, microphthalmia and primary open-angle glaucoma (Gallardo et al., 1999; Iglesias et al., 2014; Lu et al., 2019). Overexpression of *six3* and/or *six6* leads to retinal enlargement (Bernier et al., 2000; Zuber et al., 1999), rostral forebrain enlargement (Kobayashi et al., 1998), formation of ectopic retinal primordia (Loosli et al., 1999), or ectopic lens formation (Oliver et al., 1996). Thus, both loss-of-function and gain-of-function data support the conclusion that *Six3* and *Six6* are involved in formation of eye and/or brain structures at multiple developmental points in vertebrates.

*Pax6*, a member of the PAX family of TFs containing a paired-domain (PD) and a HD is essential for both lens formation and normal retina formation in human and mice (Ashery-Padan et al., 2000; Glaser et al., 1994). Similarly, *X. tropicalis pax6* null mutant embryos do not develop a lens and retinal tissue degenerates during larval stages (Nakayama et al., 2015).

As described here, both *Pax6* and *Six3* contribute to lens formation, a process in *Xenopus* that has been described as involving multiple stages, beginning at gastrulation, when presumptive lens ectoderm (PLE) is first competent to form a lens, followed by the “bias” stage when lens-inducing signals emanate from the neural plate/retina area, then determination at the neural tube stage and subsequently by differentiation, when crystallin synthesis occurs (Grainger, 1992; Grainger et al., 1997). Gene regulation studies during the lens induction process (Gunhaga, 2011; Jin et al., 2012; Ogino et al., 2012) reveal essential signaling and TF networks, e.g., in *X. tropicalis* showing that Delta/Notch signaling from the optic vesicle leads to activation in the PLE of *foxe3*, a forkhead class TF essential for lens development (Ogino et al., 2008). Other studies on signaling from the developing retina show that BMP4 functions as an optic vesicle-derived signal required for lens formation (Furuta and Hogan, 1998; Huang et al., 2015). However, BMP4 alone without the optic vesicle is not sufficient to mimic the lens inductive activity, suggesting that BMP4 regulates lens induction by synergistically acting with additional retinal factor(s) (Furuta and Hogan, 1998). Furthermore, *Pax6* is not required for this induction mediated by BMP4, and BMP4 is not required for *Pax6* expression either in the PLE or optic vesicle, suggesting that expression of *Pax6* and *Bmp4* appear to be regulated and function independently during lens induction (Furuta and Hogan, 1998).

Among the earliest of the transcriptional regulators activated in the PLE, *six3* and *pax6* are already active in *Xenopus* by the neural plate stage, the lens “bias” stage (Ogino et al., 2012; Zygar et al., 1998). Tissue recombination experiments using rat *Pax6* mutant (*rSey*) embryos reveal the tissue-autonomous requirement of *Pax6* for lens formation, at least from the neural tube stage onward, since the PLE from *rSey* embryos fails to form lenses when transplanted to WT hosts at this stage but not vice versa (Fujiwara et al., 1994).

Taken together, *Pax6* is essential but not sufficient for the PLE to become lens without having inductive signal(s) from the optic vesicle. Here, using the advantage of the *Xenopus six3* mutant to continuously follow all early events of eye formation, we show that Six3 is essential for lens formation, acting independently of *pax6* early in development, and that its function, is, in significant part, mediated non-autonomously, through its activity in the PR, in regulating BMP and Notch signaling.

## RESULTS

### The relatively mild phenotype of the *Xenopus six3* mutant allows us to investigate its role during all stages of eye development

We have previously reported F0 mutations in the *six3* locus in *X. tropicalis* by CRISPR/Cas9 gene editing (Nakayama et al., 2013). As illustrated in Fig.1A, there are two characteristic domains, the SIX domain (SD) and homeodomain (HD), both of which are 100% identical at the amino-acid level between *X. tropicalis* and human SIX3. The mutant used in the current studies results in embryonic lethality and has a premature termination mutation with a truncated open reading frame (ORF) of only 12 amino acids (aa) into the SD. A study of human *SIX3* mutations found in HPE patients indicates that mutated SIX3 proteins could have some activity only if 40 aa or more of the SD are intact (Domené et al., 2008) and we therefore reason that our mutant is nonfunctional. Other *Xenopus six3* mutant lines indicate that the “LEET” amino acid sequence in the SD is essential for *six3* function (Fig. S1).

Compared to wildtype (WT) siblings (either +/+ embryos or +/-embryos, which show no phenotype) (Fig.1C, C’), mutants (-/-) (Fig.1D, D’) have a reduced forebrain, a fused nasal pit, and smaller eyes with defects that vary in severity from one embryo to another as described more in detail in the next section, and as seen with human HPE phenotype. However, as opposed to the eyeless phenotype of null mice (Lagutin et al., 2003), the *Xenopus* mutant phenotype is less severe. Repression of Wnt signaling by Six3 has been reported to be essential for the formation of the anterior forebrain structures in mouse (Lagutin et al., 2003). The milder phenotype observed in the frog we reason is likely due to the lack of change in expression domain of *wnt1* during the neural plate stage (data not shown). This apparent difference in regulation has fortuitously allowed us to investigate the role of *six3* during all stages of lens and eye development.

In addition, we examined the functional relationship of *six3* and *six6* in *Xenopus*, since they have overlapping expression domains (Ghanbari et al., 2001), and additive effects between Six3 and Six6 during murine retina development was reported (Diacou et al., 2018). To examine this relationship in *Xenopus* we first assayed *six6* expression in the *six3* mutant by whole-mount *in situ* hybridization (WISH), but we did not observe reduction in *six6* expression (data not shown). We also generated mutations in the *six6* locus (Fig. 1B), targeting near the human *SIX6* missense HD mutation T165S found in anophthalmia and/or microphthalmia patients (Gallardo et al., 2004). A germline mutation of *six6* (Fig.1B), where the resultant Six6 protein is truncated at 159 aa followed by unrelated 7 aa caused by a frameshift has no apparent phenotype and this homozygous *six6* mutant can survive to adulthood. Analogous to the above mentioned human *SIX3* mutations analysis (Domené et al., 2008), the frog *six6* mutation, we postulate, could be hypomorphic due to the presence of an intact SD. We tested for complementary activity with *six3* by generating stable *six3* and *six6* double mutants and investigating the phenotypes in this double mutant line. Upon crossing heterozygous double mutants, i.e., *six3* (+/-), *six6* (+/-), we observed five typical phenotypes in offspring at st.45-46 (all stages according to Nieuwkoop and Faber, 1967): the first two are indistinguishable from either WT (not shown, as seen in Fig.1C, C’) or *six3* mutant denoted as type I phenotype (as seen in Fig.1D, D’). The other two phenotypes result in a tiny eye (Fig.1E, E’), denoted as type III, or an extremely rudimentary eye (Fig.1F, F’), denoted as type IV. Some embryos showed intermediate eye phenotypes between type I and type III, denoted as type II (not shown). Type III and IV mutants were always either *six3* mutants carrying single or double mutant copies of mutant *six6*, respectively. In summary, while the *Xenopus* data shows that the additional *six6* mutation accentuates the *Xenopus six3* phenotype, in the *six3* mutant, *six6* is not normally affected as it is in the mouse *Six3* mutant, likely because, overall, the mouse *Six3* mutation has a stronger impact on head and eye development, through its effect on *Wnt1* and *Six6*, neither of which is seen in the *X. tropicalis six3* mutant.

**Fig. 1.**
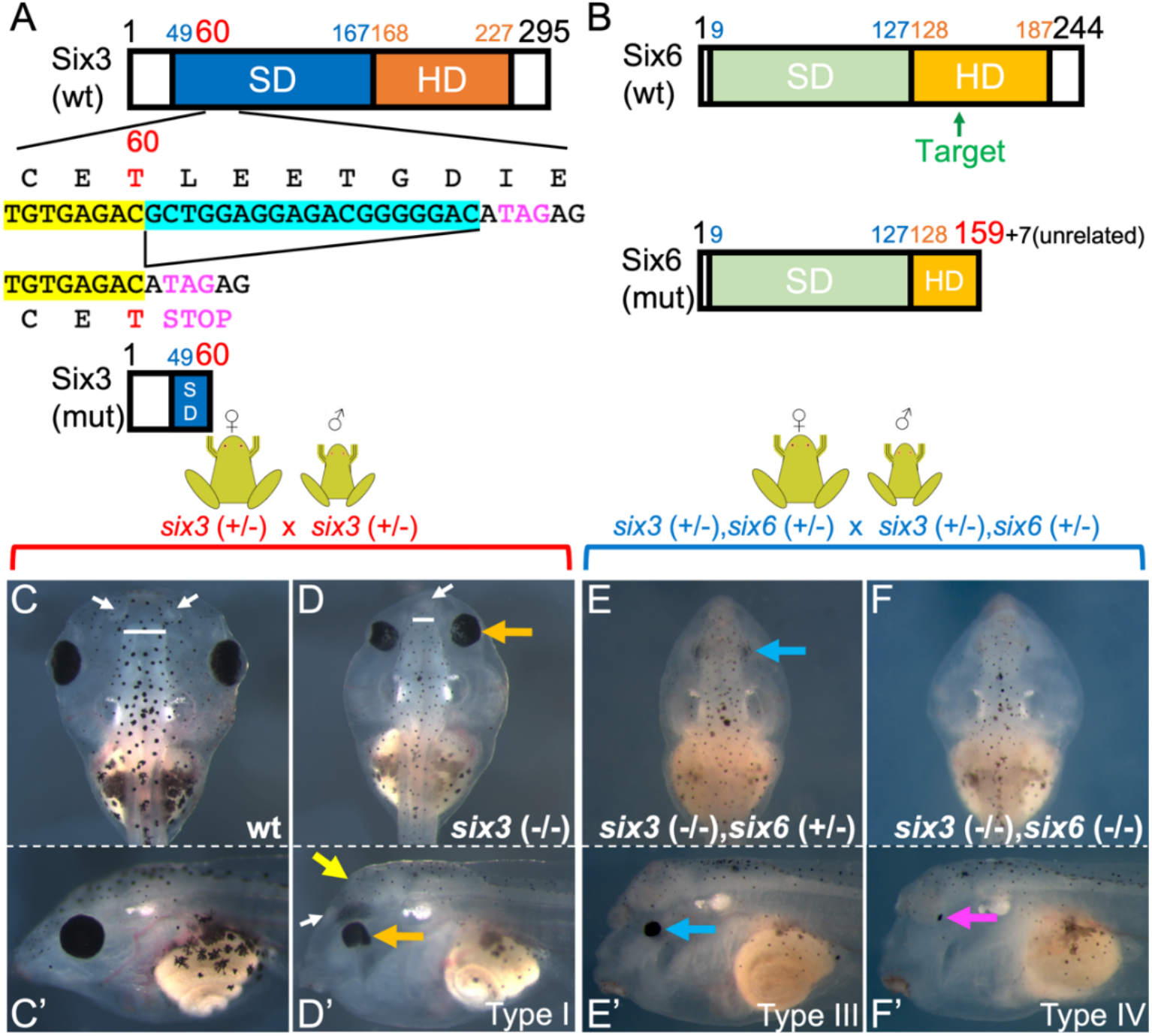
Homozygous *six3* mutation in *X. tropicalis* causes holoprosencephaly and eye defects, leading to loss of the eye by additional mutation of *six6*. (A) Diagram of the *six3* mutation. SD, SIX domain (amino acids 49-167) shown in blue. HD, homeodomain (amino acids 168-227) shown in dark orange. A cyan-highlighted region indicates deleted sequence in the mutant locus, resulting in a premature stop codon (TAG shown in magenta). The resultant protein ends at amino acid 60 (T in red). A yellow-highlighted portion indicates the 5’ junction of the deletion. (B) Diagram of the *six6* mutation (CRISPR target position shown by green arrow). SD, SIX domain (amino acids 9-127) shown in light green. HD, WT homeodomain (amino acids 128-187) and truncated homeodomain (amino acids 128-159) in the mutant shown in light orange. The mutant partial HD is followed by 7 unrelated amino acids caused by a frameshift in this mutation. A WT st.45/46 tadpole, dorsal (C) and lateral (C’) views. Representative phenotype of a homozygous *six3* mutant tadpole, dorsal (D) and lateral (D’) views. White lines in (C) and (D) denote the width of the anterior edges of the telencephalon, showing the narrower telencephalon in the mutant. White arrows (C, D, D’) show the nasal pits, which are fused in the mutant. Orange arrows (D, D’) show abnormal eyes in the mutant. Yellow arrow denotes the reduced telencephalon. Typical phenotype of *six3* (-/-), *six6* (+/-) mutant, dorsal (E) and lateral (E’) views. Blue arrows (E, E’) indicate eyes with severer defects than is seen in *six3* (-/-) mutants (D, D’, orange arrows). Typical eyeless phenotype of the *six3* (-/-), *six6* (-/-) double mutant, dorsal (F) and lateral (F’) views. Magenta arrow (F’) indicates the eyeless phenotype with a residual amount of retinal pigment epithelium (RPE) pigment. All embryos at st.45/46.

### The *Xenopus six3* mutant reveals a previously unrecognized essential role for *six3* in retina and lens formation

The eye defects seen in the *six3* mutant can be grouped into three types based on phenotype severity, scored from “+” (mild) to “+++” (most severe) with “-” for WT based on histology (Fig. 2A). In WT embryos at st.45, the lens is highly differentiated, consisting of a clear core of fiber cells (red arrow) surrounded by epithelial lens cells (orange arrow) and well-organized, laminated retina (green arrow) (Fig. 2A, wild type). In mutants (Fig. 2A, *six3* mutant), by contrast, nearly half of the mutant eyes scored (43.75%, 7 eyes out of 16 eyes examined from 8 embryos) did not have a lens and the retina was disorganized without having obvious layers (scored +++). Five eyes (31.25%) had a small lens-like structure consisting of only epithelial cells together with relatively organized (data not shown) or sometimes even disorganized retina (++). By contrast, four scored eyes (25%) had relatively well-formed but still abnormal retinas and a small lens consisting of partially differentiated fiber (white arrow) and epithelial cells (scored as +). In an individual mutant embryo, both eyes tended to show the same phenotype severity. We further noted that mutant embryos having more organized retinas seem to have more differentiated lenses.

**Fig. 2.**
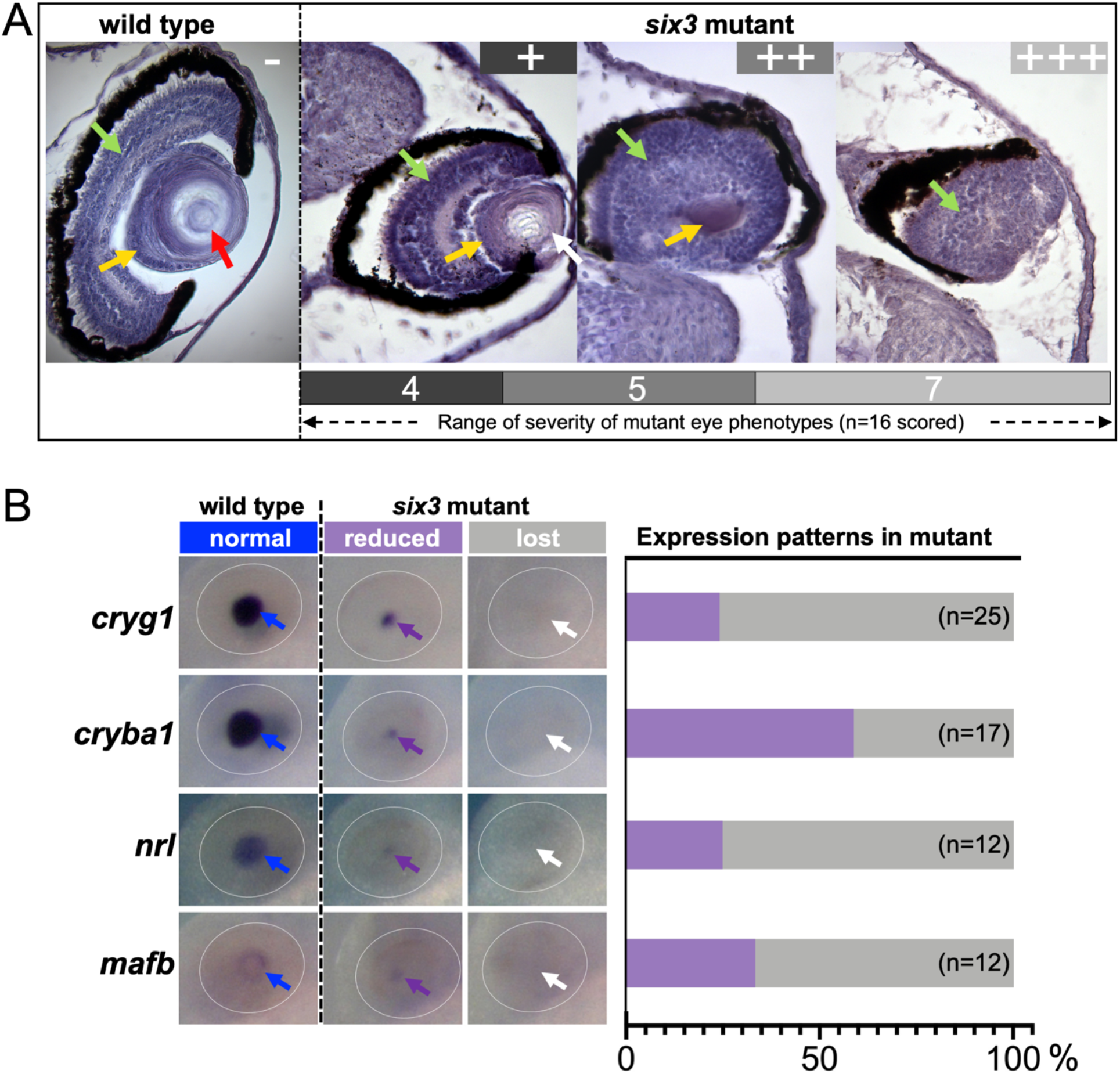
*six3* mutant shows variations in eye phenotypes. (A) Histology of eyes of WT and mutant at st.45. Green arrows, retinal regions. Orange arrows, lens epithelial cells. Red arrows, well-differentiated lens fiber cells. White arrows partially differentiated lens fiber cells. The severity of *six3* mutant eye phenotypes were scored from + (dark gray), ++ (gray), and +++ (light gray) as described in the text. The numbers of eyes scored are shown in the bar under mutant histology. (B) Expression profiles of representative markers expressed in the lens at st.37/38 in WT and mutant embryos. The tested markers are shown on the left side of images. Left-most column images show expression in WT. The right two columns show expression patterns in the mutant: reduced level of expression (purple) and no expression (gray). Dotted ovals are retina regions. Arrows point toward lens regions. The percentage of each pattern is shown in the bar graph at the right. The number of eyes examined is shown in parentheses.

Owing to the limitations in performing quantitative assessments by histology, we examined representative markers expressed during lens development at st.37/38, by when, for example, a lens fiber marker, *cryg1*, is highly expressed in WT embryos (e.g., Jin et al., 2012). Consistent with the histology results, we observed by WISH that *cryg1* expression is completely lost in the majority (76%) of mutant embryos examined, whereas 24% of mutant embryos had severely reduced levels of *cryg1* expression (Fig. 2B, *cryg1*). This correlated well with the histology result that 25% had lenses with partially differentiated fiber cells but the remaining mutant eyes (75%, “++” and “+++” together) examined did not have obvious fiber cells. Another marker *cryba1*, whose expression is known to be the earliest among crystallin genes, first activated in the lens placode (Day and Beck, 2011), was still seen in 59% of mutant embryos, though at a severely reduced level and in a smaller domain compared to WT embryos (Fig. 2B, *cryba1*, reduced). This is consistent with histology data where 56% eyes examined (i.e., “+” and “++” together) had a small lens or lentoid. In summary, both histology and expression of lens markers imply that in the *six3* mutants about half of their eyes fail to form lenses and the other half have very small lenses or a primitive lentoid structure.

The members of the large Maf family of basic leucine zipper lens TF’s function in proliferation of epithelial cells or the differentiation of fiber cells, and to regulate transcription of crystallin genes (Ishibashi and Yasuda, 2001; Ogino and Yasuda, 2000). In *Xenopus* (Coolen et al., 2005; Ishibashi and Yasuda, 2001), *mafb* is the earliest large Maf gene expressed in the PLE, as early as st.22, whereas *nrl* is activated slightly later, at st.24. Eventually *mafb* transcripts are seen in lens epithelium and *nrl* in fiber cells, and expression of both persists at least until st.37/38 in WT embryos (Fig. 2B, WT). In *six3* mutants, expression of both is largely lost (66.7% and 75% for *mafb* and *nrl*, respectively) (Fig. 2B). Taken together these results show serious defects in both the retina and lens. In some *six3* mutant embryos a degree of lens differentiation occurs. These results led us to investigate if there are intrinsic defects in the lens and/or whether non-cell autonomous signaling could be impacting lens formation in the mutants.

### *pax6* expression is independent of *six3* in the *Xenopus eye*

Focusing on the lens phenotype observed in the *six3* mutant (Fig. 2) we set out to determine if the phenotype observed is due to impact of *six3* alone or if *pax6*, an essential gene for lens formation (Ashery-Padan et al., 2000), is also disrupted. Since several reports in the literature have suggested the existence of mutual regulation between *Pax6* and *Six3* (Ashery-Padan et al., 2000; Goudreau et al., 2002; Liu et al., 2006) we investigated whether this mechanism is involved in the phenotypes seen in *Xenopus*. Following up the observations in previous sections, we examined younger embryos, at st.15, the lens-forming “bias” stage, for expression of *six3* by WISH in the *six3* mutant as well as in the *pax6* mutant that does not form lens (Nakayama et al., 2015). In both mutants (Fig. 3A’, B’) the expression patterns of *six3* in the PLE (blue arrows) and/or presumptive retina (PR) region (white arrows) are indistinguishable from WT (Fig. 3A, B). We also examined *pax6* expression in the *six3* mutant (Fig. 3C, C’, D, D’) observing indistinguishable expression patterns in the PLE (orange arrows) and PR (white arrows) between mutant and WT during st.15-18. These results indicate that during these stages, *six3* and *pax6* expression are not mutually regulated in the PLE of *Xenopus*. We also investigated *pax6* expression in the *six3* mutant at st.24/25 by WISH (Fig. S2 A, A’) where its expression appears unchanged in both in the PR and the PLE between WT and *six3* mutant embryos as revealed by histology (Fig. S2 B, B’). The presence of *pax6* expression throughout the lens-forming process, we reason, could be a factor that led to the presence of lens like structures in some *six3* mutant animals. This is further supported by expression of lens markers like *nrl* which are regulated by Pax6 activity (Reza et al., 2002).

**Fig. 3.**
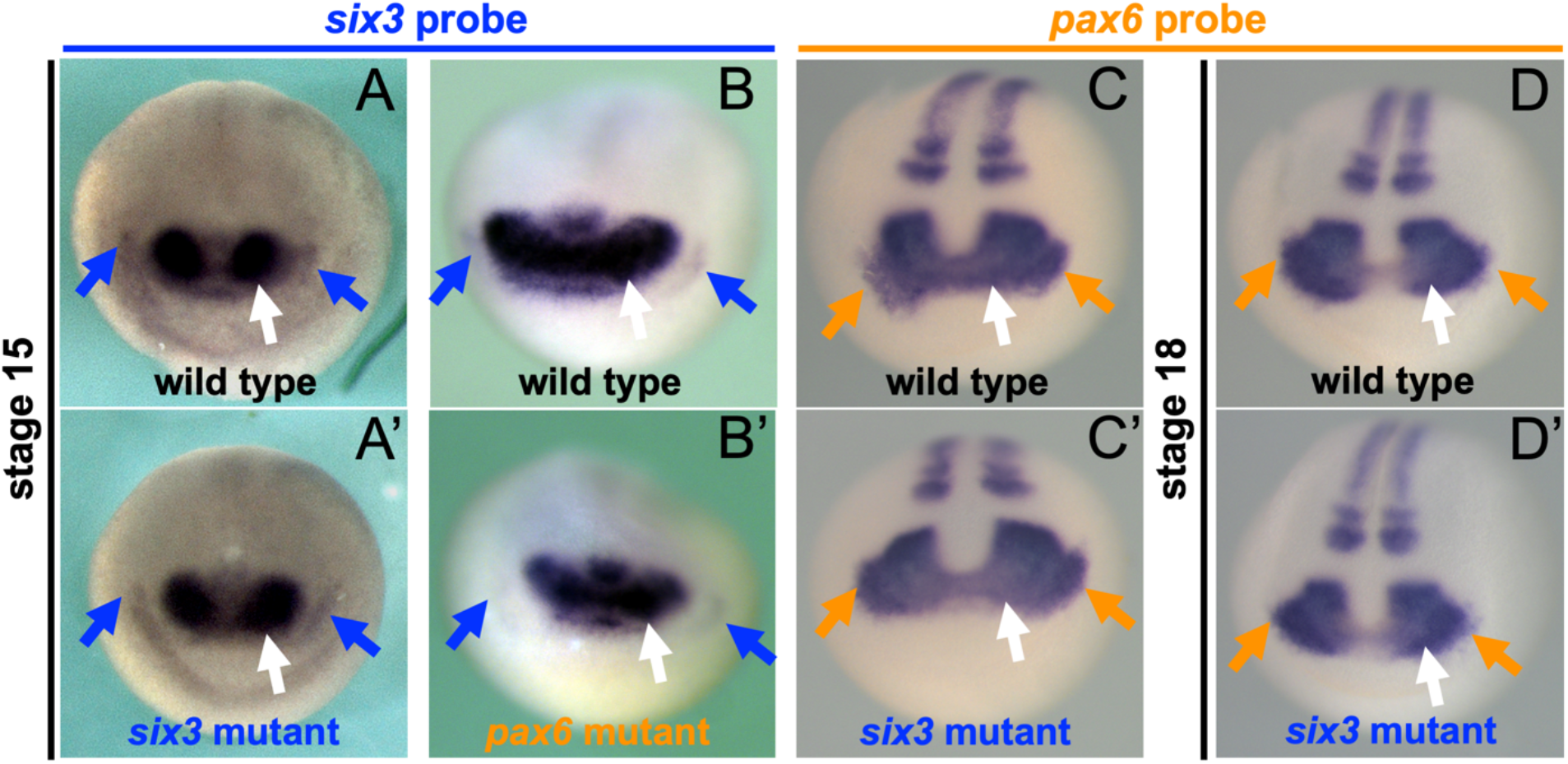
Expression of *pax6* and *six3* are not perturbed in *six3* and *pax6* mutant respectively. (A, A’), (B, B’) *six3* expression in WT (A, B), *six3* mutant (A’) and *pax6* mutant (B’) at st.15. (C, C’), (D, D’) *pax6* expression to detect *pax6* in WT (C, D) and *six3* mutant (C’, D’) at st.15 (C, C’) and st.18 (D, D’). Blue, orange arrows indicate PLE region. White arrows indicate the PR region.

Results described here show that phenotypes in the *six3* mutant are primarily due to the strong effect of *six3* during early stages of lens and retina formation even in the presence of normal expression of *pax6*. We also show that *pax6* expression is independent of Six3 activity, implying that *six3* plays a key, and independent role from *pax6* during lens formation.

### Failure of lens determination is primarily non-autonomous in the *six3* mutant

To test whether an intrinsic defect in the PLE or a defect of PR, or both, in the *six3* mutant leads to lens formation failure, we performed transplant experiments using st.15, bias-stage, embryos as illustrated in Fig. 4A. Five independent experiments used offspring obtained from heterozygous mutant crosses for both hosts and donors, and one experiment used WT hosts to increase the probability of obtaining mutant donor/wild type hosts (mut/wt). A total of 65 successful transplants resulted in 35 wt/wt, 7 mut/wt, 18 wt/mut (and 5 mut/mut) cases, from each of which randomly selected embryos were assayed for *cryg1* expression at st.37/38 (Fig. 4B). A positive control experiment (Fig. 4B, wt/wt) showed 100% of the recombined embryos examined (N=12) had indistinguishable levels (++++) of expression of *cryg1* compared to the unoperated side. If a PLE defect in the *six3* mutant were the cause of loss of the lens, recombinants of mutant PLE and WT host (mut/wt) would not form lenses, but the result was not consistent with this prediction:100% recombined embryos tested (N=5) had indistinguishable level (++++) of expression of *cryg1* from that seen in the unoperated side (Fig. 4B, mut/wt). This clearly suggests that *six3* is not necessary in the PLE to form the lens. By contrast, WT PLE combined with mutant host (wt/mut) showed either complete loss of *cryg1* expression (∼60%, 7 out of 12) or drastically reduced level of *cryg1* expression (∼40%, 5 out of 12) in recombined embryos, implying that a PR defect is implicated in the loss of the *six3* mutant lens.

**Fig. 4.**
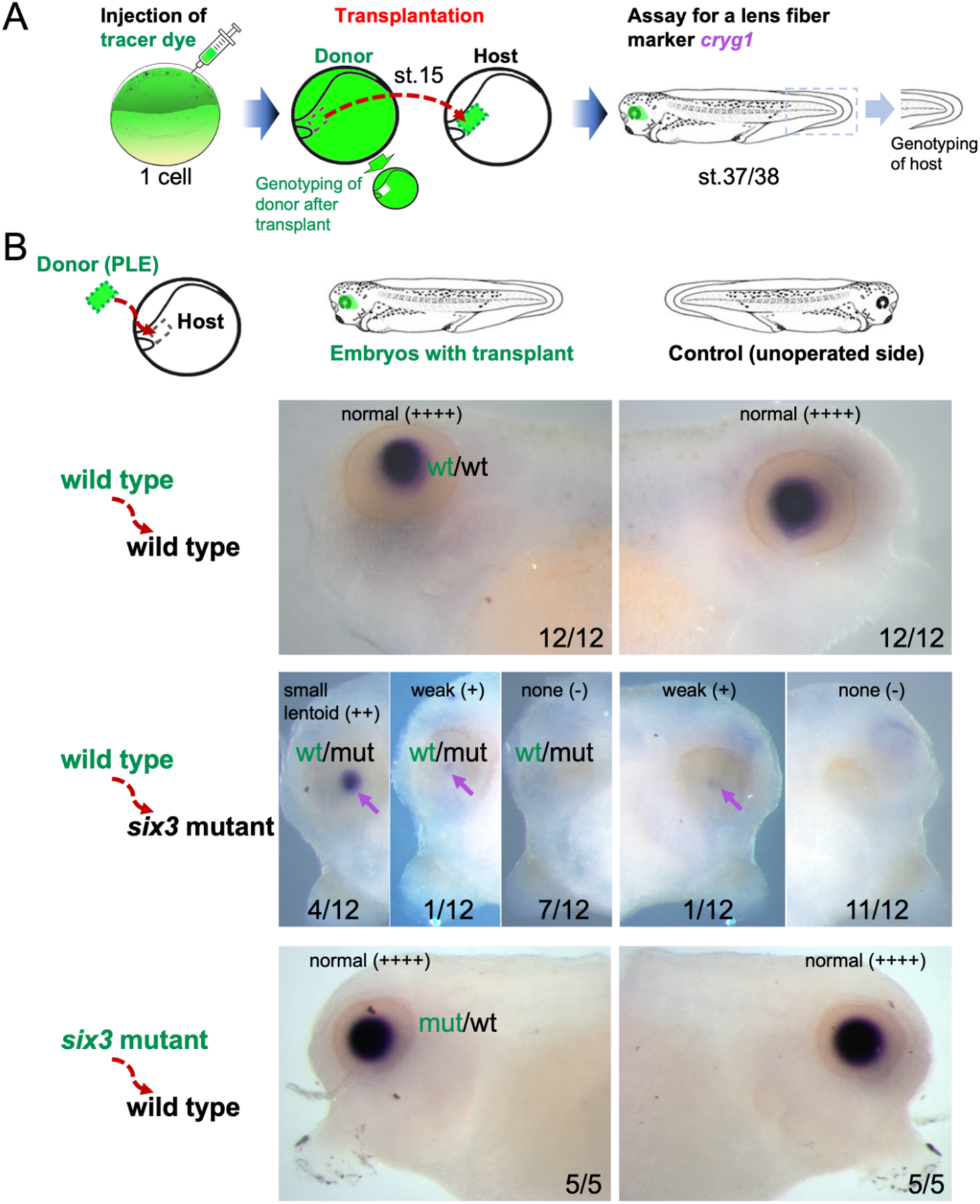
Transplanted WT PLE in *six3* mutant host fails to make lens. (A) Diagram of experimental procedure. (B) WISH to detect *cryg1*. On the left are schematics to show combinations of the PLE (shown in green) and host (in black). The right-side panels show images of representative staining of *cryg1* for each transplanted side and unoperated side as illustrated above the panels. normal (++++), WT level of *cryg1*. small lentoid (++), weak (+), none (–), show progressively reduced levels of *cryg1*.

These results support the hypothesis that failure of lens formation in *six3* mutants occurs mainly non-cell autonomously due to failure in PR signaling. However, since ∼40% wt/mut recombinants had weak expression of *cryg1*, of marginal statistical difference (*p*-value = 0.06 by two-tailed Mann–Whitney U test) compared to the control side, WT PLE may also have ability to mount some response to recover from the loss of signaling from the mutant PR. We, therefore, would not rule out that *six3* also has a degree of cell autonomous function in the PLE for lens formation, which remains for future study. However, our transplant data indicate that the lens defect observed in the *six3* mutant is largely a non-cell autonomous defect during lens induction.

### Notch signaling components and *foxe3* expression are downregulated in the *six3* mutant

To investigate the potential role of optic vesicle signaling in inducing lens formation that is disrupted in the *six3* mutant, we first focused on a previously identified process in *Xenopus*, that activation of *foxe3*, a forkhead TF gene essential for lens formation in *Xenopus*, is mediated by Notch signaling from the optic vesicle (Ogino et al., 2008). The *foxe3* gene is known to be expressed from the early neural plate stage, followed by expression primarily in the PLE overlying the optic vesicle when commitment to the lens fate occurs during neural tube stages. Its expression is downregulated as committed lens cells undergo terminal differentiation, and from st.30 onward its expression is mainly seen only in the anterior epithelial cells of the lens vesicle (Kenyon et al., 1999). By st.21, the expression becomes consistently strong in WT embryos (Fig. 5A), whereas in the mutant the expression is reduced compared to WT (Fig. 5A’). At st.24/25, *foxe3* expression was no longer detectable by WISH in the mutant (Fig. 5B’).

**Fig. 5.**
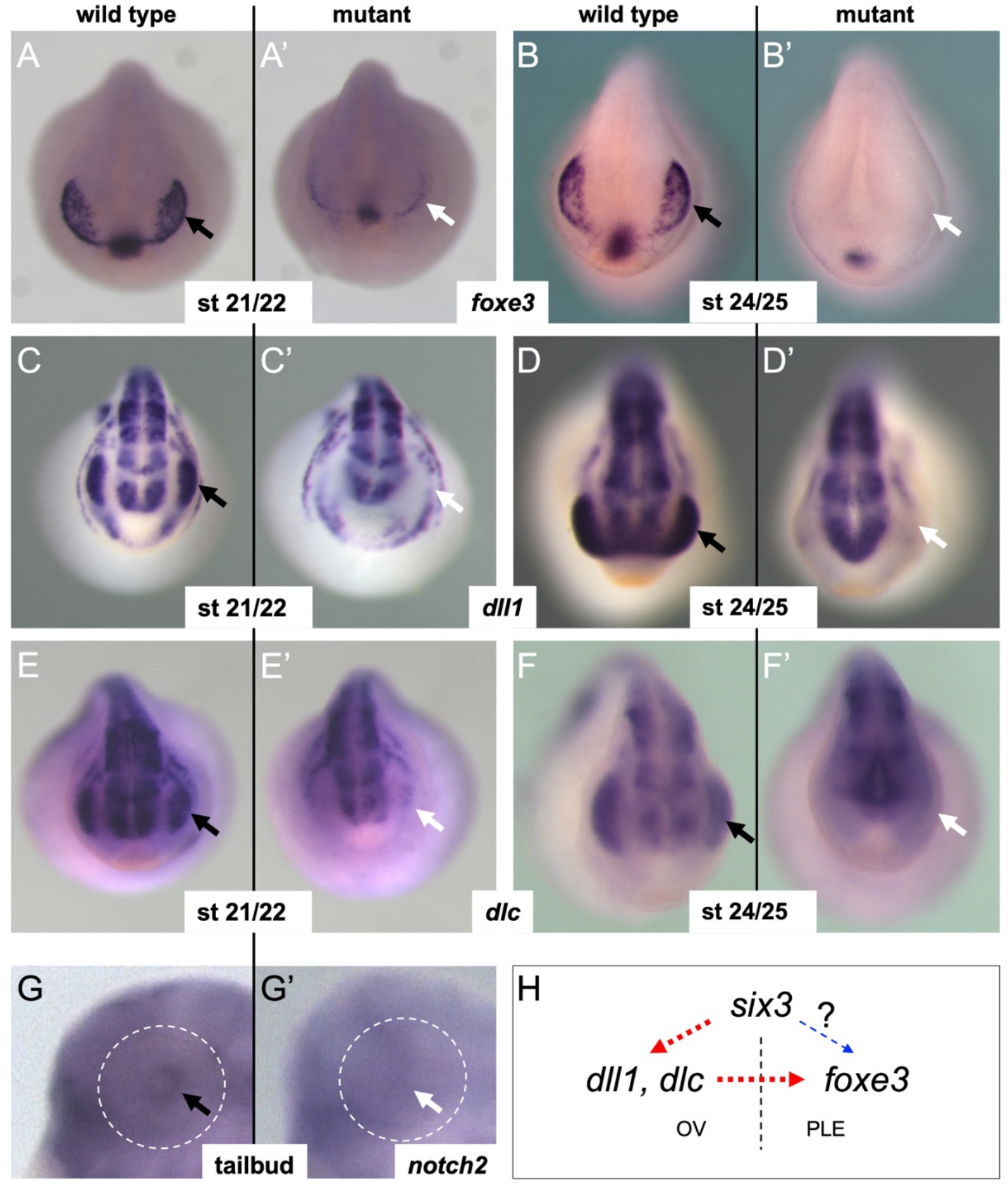
*foxe3* and its known upstream Notch signaling components are downregulated in the *six3* mutant. (A-B’) *foxe3* expression at st.21/22 (A, A’) and st.24/25 (B, B’) in WT (A, B) and *six3* mutant (A’, B’). (C-D’) *dll1* expression at st.21/22 (C, C’) and st.24/25 (D, D’) in WT (C, D) and *six3* mutant (C’, D’). (E-F’) *dlc* expression at st.21/22 (E, E’) and st.24/25 (F, F’) in WT (E, F) and *six3* mutant (E’, F’). (G) *notch2* expression in WT and in *six3* mutant (G’) at st.32. Dotted circles indicate the retina regions, arrows point to the lens. (H) Possible gene hierarchy to regulate *foxe3*. See the main text for details. OV, optic vesicle. PLE, presumptive lens ectoderm. Black arrows, expression in the eye region in WT. White arrows, expression in the *six3* mutant eye region.

The Notch signaling described above is mediated by Dll1 and/or Dlc ligands expressed in the underlying PR, activating the Notch2 receptor in the PLE, leading to activation of *foxe3* (Ogino et al., 2008). We therefore examined *dll1* and *dlc* expression at st.21 and st.24 and found that both *dll1* and *dlc* are lost in the *six3* mutant (Fig. 5C’-F’). *notch2* is expressed in a broad region including the retina at these stages (Ogino et al., 2008), though we could not conclusively determine whether its PLE expression is perturbed in the *six3* mutant at st.21-24 because of low and variable level of expression by WISH (data not shown), but its expression was clearly lost at st.32 (Fig. 5G, G’). In Summary, data presented here indicate that *six3* functions, at least in part, upstream of *dll1* and/or *dlc* in the PR, thereby activating *foxe3* in the PLE (Fig. 5H, red dotted arrows).

### BMP signaling pathway is perturbed in the *six3* mutant

In addition to the role of Notch signaling in lens formation, it has been shown in mice that BMP4 and BMP7 are essential for lens induction (Dudley et al., 1995; Furuta and Hogan, 1998; Huang et al., 2015; Luo et al., 1995). These findings motivated us to examine expression of *bmp4* and *bmp7*.*1* (the *Xenopus* ortholog of mouse *Bmp7*) in *X. tropicalis* at st.21, when we have shown above that defects in gene expression in the PR, potentially leading to loss of the lens, have already occurred in the *six3* mutant.

WISH shows that both BMP genes are significantly reduced in the *six3* mutant at st.21 (Fig. 6A-B’). To assay the functional consequence of this, we analyzed levels of phosphorylated P-Smad1/9, which provides a direct readout of BMP pathway activity (Faure et al., 2000), at st.24 (Fig. 6D-H). The data shows that P-Smad1/9 levels are reduced in the PLE of the mutant but not necessarily reduced in the PR underlying the PLE of the mutant compared to the levels in WT.

**Fig. 6.**
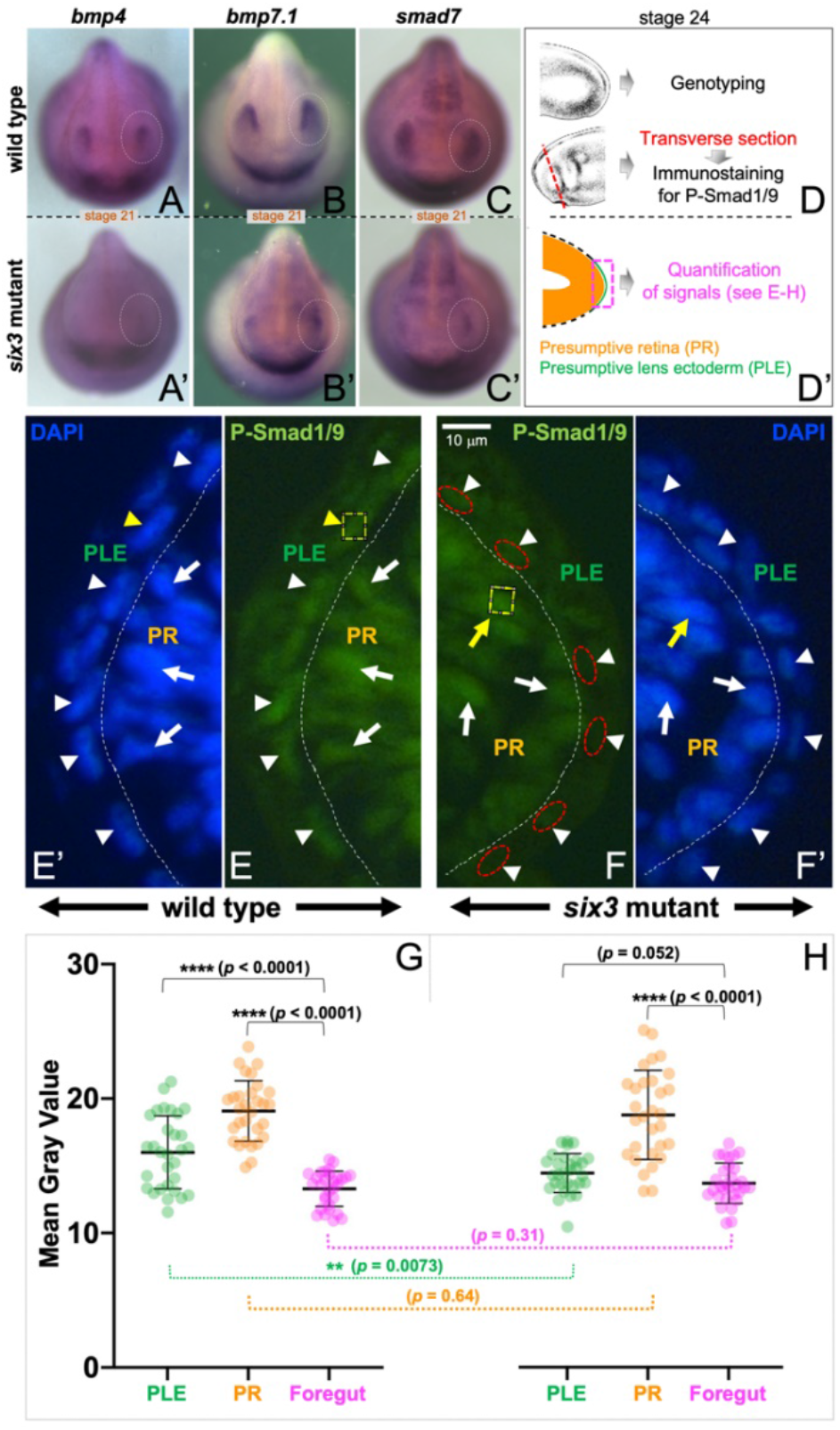
BMP signaling is perturbed in the *six3* mutant. (A-C’) Expression of *bmp4* (A, A’), *bmp7*.1 (B, B’), and *smad7* (C, C’) in WT (A, B, C) and *six3* mutant (A’, B’, C’) at st.21. Dotted white ovals indicate the eye region. (D, D’) Schematic showing the experimental procedure for P-Smad1/9 detection. The magenta box in (D’) surrounds a part of a transverse section corresponding to images shown in (E-F’) through the approximate center of the optic vesicle as indicated by the dashed red line in D. (E, F) Images of P-Smad1/9 level detected by immunofluorescence staining of WT (E) and *six3* mutant (F) embryos. Corresponding DAPI staining is in (E’, F’). White dotted lines show boundary between the PR and the PLE which are distinguishable by orientation of the cells. Arrows and arrowheads respectively indicate examples of P-Smad1/9 immunofluorescence staining (E, F) or DAPI staining (E’, F’) in the PR and PLE, among which yellow-colored arrow and arrowhead show examples of regions (dotted-yellow squares) used for calculation of Mean Gray Value shown in the graphs (G, H). Background level was measured in the pharyngeal endoderm region of the foregut (not shown). Dotted red ovals indicate examples of loss of P-Smad1/9 immunofluorescence staining (F). The scale shown in (F) also applies to (E, E’, and F’). (G, H) The data were collected and pooled from each of 3 mutant and WT sibling embryos: 10 measurements per region, i.e., total 30 Mean Gray Values for each region (for each WT and mutant) were obtained and tested by unpaired t test with Welch’s correction, two-tailed. The bars and error bars in graphs are mean ± SD (mutant PLE,14.4 ± 1.45; mutant PR, 18.7 ± 3.31; mutant foregut, 13.7 ± 1.51; WT PLE, 16.0 ± 2.72; WT PR, 19.1 ± 2.25; WT foregut, 13.3 ± 1.32. Two asterisks (**) for *p-*value < 0.01, four asterisks (****) for *p-*value < 0.0001.

To complement the studies above, we also examined *smad7* as another assay for BMP activity, since in *Xenopus* Smad7, an inhibitory Smad, is shown to be a downstream target and a component of BMP signaling to fine-tune BMP activity in a negative feedback loop (Christian and Nakayama, 1999; Nakayama et al., 1998; Nakayama et al., 2000). It is also worth mentioning that in mice, loss of SMAD7 disrupted lens differentiation and SMAD7 is implicated in lens differentiation but is not required for the induction of the lens placode (Zhang et al., 2013). By WISH we saw significant reduction of *smad7* in the *six3* mutant consistent with reductions of *bmp4* and *bmp7*.*1* (Fig. 6C, C’). Taken together, we confirmed the reduction of BMP activity in the eye region of the mutant and at the cellular level the PLE has reduced level of BMP activity in the mutant, consistent with the proposal that the loss of *bmp4* (and/or *bmp7*.*1*) expression in the PR affects the responding PLE, contributing to the loss of lens in the *six3* mutant.

Further studies reveal that this down-regulation of the BMP pathway is mediated by alterations nuclear protein genes *mab21l1* and *mab21l2*, described below, followed by a discussion of their role in regulation of BMP signaling originating in the PR.

### Additional Targets of Six3: nuclear proteins *mab21l1* and *mab21l2*

Mutations in the nuclear protein genes *mab21l1* and *mab21l2* reveal that they are both important for lens formation in mice (Yamada et al., 2003; Yamada et al., 2004) and during the stages when the lens is becoming determined they are both expressed in the lens and retina in *Xenopus* (Fig. 7), though the detailed pattern of expression differs for the two genes. Because the genes are highly conserved (93% amino acid conservation) including key structural similarities (de Oliveira Mann et al., 2016), they likely have overlapping or redundant function. Here, we focus on their expression and potential role in regulation of lens-inducing signaling within the retina, particularly in conjunction with their regulation by *six3*. As can be seen in Fig. 7, focusing on retina expression of these genes, at early stages in the PR expression of *mab2ll2* is severely reduced (st.15 and st.18; Fig. 7D/D’ and 7E/E’, white arrows) but *mab21l1* is only modestly affected (st.15 and st.18; Fig. 7A/A’ and 7B/B’, white arrows) in the *six3* mutant while at st.21 the situation is reversed, i.e., PR expression of *mab21l1* is severely impacted (Fig. 7C/C’) but *mab21l2* expression is partially restored (Fig. 7F/F’). Overall, at each of the stages one of the genes is strongly reduced in the PR, and because of their high degree of conservation, there is very likely overall reduced activity in the PR at all stages, the implication of which for lens induction is discussed in the next section. As can be seen in Fig. 7A-F’ (colored arrows), lens expression of these genes is largely unaffected at these stages apart from loss of *mab21l1* at st.21 in the *six3* mutant. Overall, there is clearly a reduction of *mab21l1/mab21l2* in the PR during lens determination and, as discussed below, this gene activity plays a key role in lens induction. While other studies show that *Pax6* mutations also affect at least *mab21l1* expression, in both mice (Yamada et al., 2003) and *X. tropicalis* (Nakayama et al., 2015), this finding amplifies what was proposed earlier: that there is a separate role for *six3* in regulating expression of these genes as shown by the data presented here.

**Fig. 7.**
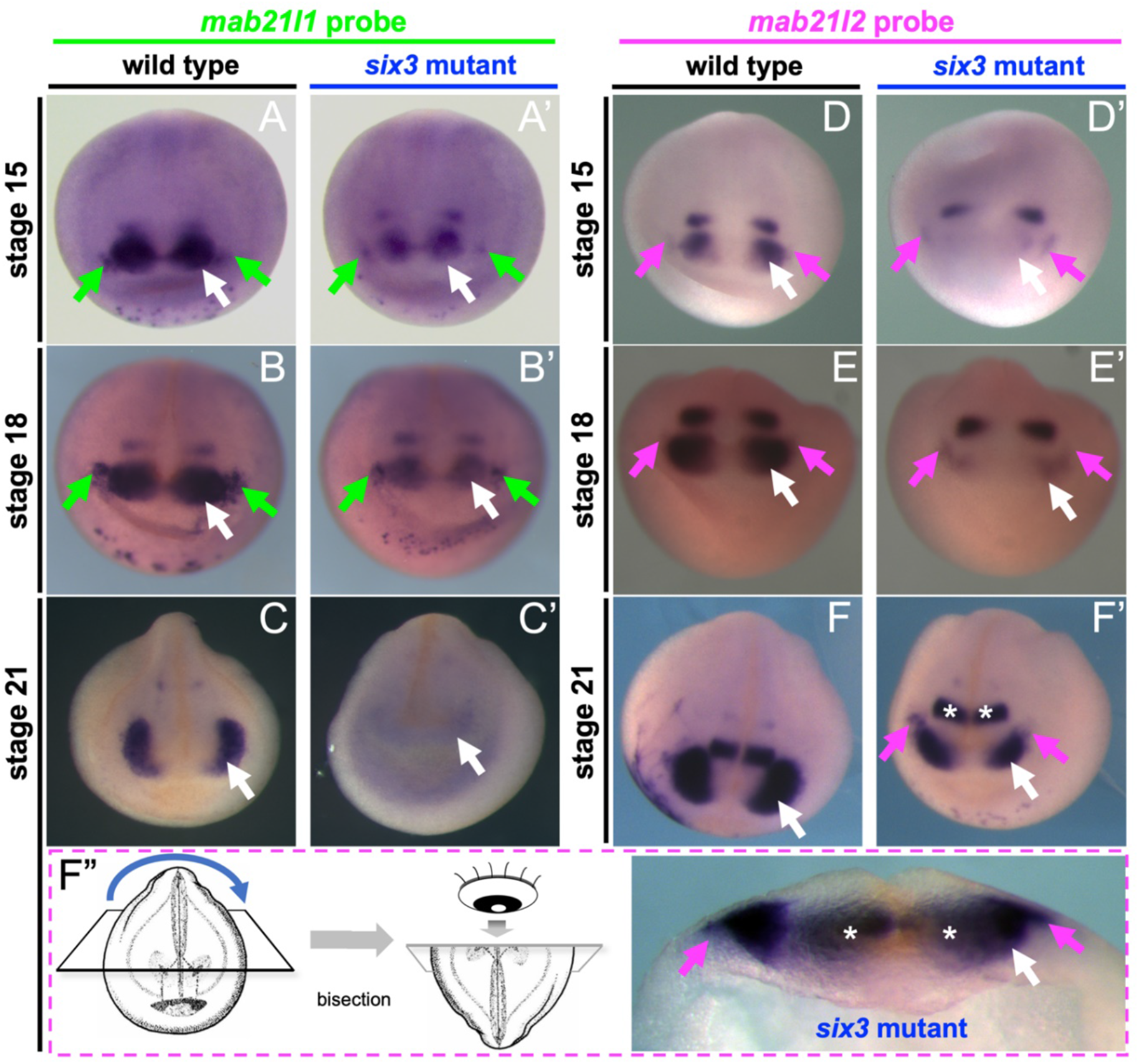
Expression of *mab21l1* and *mab21l2* in the *six3* mutant. (A, A’), (B, B’), (C, C’) WISH to detect *mab21l1* in WT (A, B, C) and *six3* mutant (A’, B’, C’) at st.15 (A, A’), st.18 (B, B’), and st.21 (C, C’). (D, D’), (E, E’), (F, F’) WISH to detect *mab21l2* in WT (D, E, F) and *six3* mutant (D’, E’, F’) at st.15 (D, D’), st.18 (E, E’), and st.21 (F, F’). (F”) Schematic of bisected embryo showing top half rotated 180 degrees as shown (blue arrow), allowing a section through the eyes and lens to be visualized as shown on the right (F’), where white asterisks indicate the midbrain region (F’, F”). Green, magenta arrows indicate PLE region. White arrows indicate the PR region. Anterior is top of image (F”. Right).

### BMP activity and lens formation is rescued in the *six3* mutant by injecting *mab21l1* mRNA

Since mis-expression of *mab21l1* and/or *mab21l2* genes and BMP signaling are apparent in the *six3* mutant at these early stages, and in addition, previous studies have shown that *mab21l2* interacts with *smad1* (Baldessari et al., 2004) and impact BMP signaling, we pursued the possible connection between alterations in *mab21l1/mab21l2* and BMP signaling in lens induction. Therefore, we tested whether injection of *mab21l1* mRNA could rescue the *six3* mutant lens phenotype, hypothesizing that this may occur via effects on BMP-mediated pathway (Fig. 8).

**Fig. 8.**
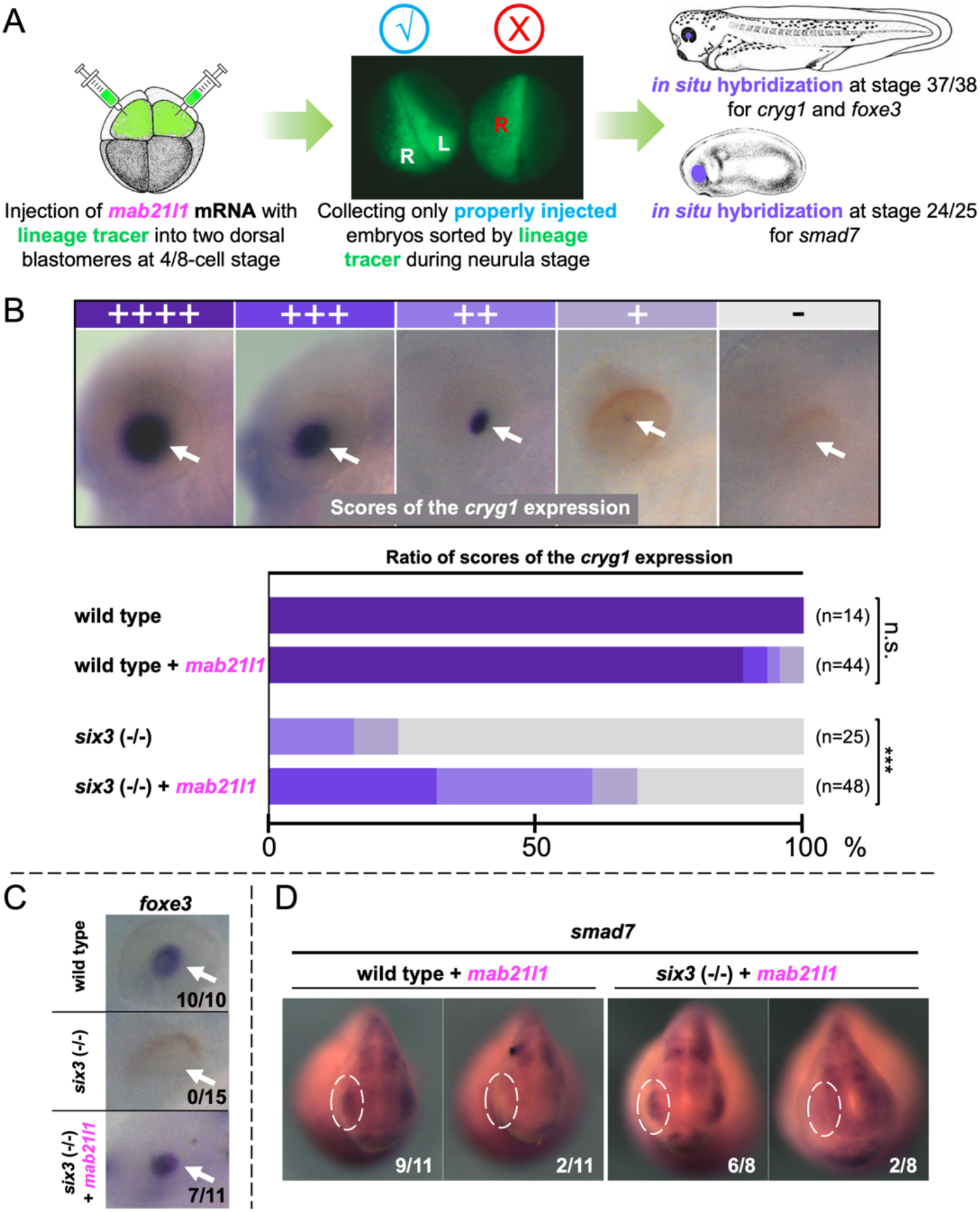
Injection of *mab21l1* mRNA partially restores expression of lens markers and BMP activity in the *six3* mutant. (A) Experimental procedure. See Method for details. (B) st.37/38 *cryg1* expression levels and their corresponding scales (++++ to -) and color codes (dark purple to pale gray) for scoring, are summarized in the graphs. n, number of eyes scored. The data set for *six3* (-/-) is the same as Fig. 2B where scores of ++ and + shown here are combined and mentioned as “reduced”. n.s., *p-*value is 0.57 and ***, *p-*value < 0.001 by two-tailed Mann-Whitney U test. (C) The representative *foxe3* expression pattern in WT, mutant, and *mab21l1*-injected mutant embryos at st.37/38. (D) The representative st.24/25 *smad7* expression pattern in WT, *mab21l1*-injected WT, mutant, and *mab21l1*-injected mutant embryos. Numbers of embryos showing indicated expression pattern (D) or positive cases (C) per numbers of embryos examined are shown.

We initially determined whether lens formation is rescued in the *six3* mutant by WISH for *cryg1* (Fig. 8A, B). We first investigated if injection of *mab21l1* mRNA causes any phenotypes by itself in WT embryos, observing some minor reduction in *cryg1* expression likely due to injection artifacts. When injecting in mutants, we observed rescue of *cryg1* expression (Fig. 8B). Although we did not see the WT level (++++) of rescued expression, 31.3% eyes showed +++ level of *cryg1* expression which was never observed in non-injected mutant eyes, and 29.2% ++ level of expression in injected mutant eyes (as opposed to 16% in non-injected mutant eyes). The percentages of + level is not different between non-injected and injected embryo eyes. Overall, only 31.3% of the injected embryos show no *cryg1* expression as opposed to 76% in non-injected embryos.

We also tested if *foxe3* is rescued by *mab21l1* injections (Fig. 8C) because its expression is reduced in the murine *mab21l1* mutant (Yamada et al., 2003). As opposed to *cryg1*, a complete loss of *foxe3* expression has already occurred by st.24 in the *six3* mutant (Fig. 5B’), and the phenotype is not variable in the mutant thereafter. Seven out of 11 injected mutant embryos examined showed rescue of *foxe3* expression in both eyes of 6 embryos and in one eye of one embryo.

We next examined if BMP activity could be rescued by *mab21l1* mRNA injection (Fig. 8D). Here, we looked at *smad7* expression, as discussed above, as a readout of activation of BMP pathway. Similar to *cryg1* described above, some WT embryos (2 out of 11) had reduction of *smad7* upon injection, never seen in non-injected embryos, which we attribute to a low-level injection artifact. Six embryos out of eight injected mutant embryos showed rescue of *smad7* expression at a level indistinguishable from WT. Two negative cases could also be due to injection artifacts after rescue. In summary, we cannot statistically distinguish WT embryos from *six3* mutant embryos injected with *mab21l1* (*p*-value is 0.72 by a chi-square test) for the level of *smad7* expression. Taken together, these results clearly show that a significant fraction of mutant lens phenotypes were rescued by *mab21l1* mRNA injection which we argue is at least in part due to rescue of BMP signaling.

## DISCUSSION

This study presents novel and underappreciated roles for *six3* in lens formation, using a *Xenopus* mutant, which allows access to eye development at all stages, contrasting the mouse *Six3* mutant in this regard. We show that this activity of *six3* is mediated in large part non-cell autonomously from the PR and is independent of *pax6* activity in early development. Fig. 9 summarizes the key findings presented here, primarily the impact of *six3* on BMP signaling through effects on *bmp4, bmp7*.*1*, and *smad7* that we propose are mediated by *mab21l1* and *mab21l2*, as well as on Notch signaling (through effects on *dll1* and *dlc*) in addition to other as yet unidentified factor(s) “X”. Importantly, another example of non-cell autonomous induction of the lens via overexpressed Six3 without “direct” induction of *pax6* was reported in medaka fish, which occurred only nearby the otic vesicle (Oliver et al., 1996). They speculated that Six3 misexpression may induce a soluble factor that changed the bias of the otic placode towards a lens fate. Since the otic vesicle is expressing *bmp4* naturally (e.g., Chatterjee et al., 2010), this may be coincidentally mimicking the native situation of the PLE and optic vesicle. Also, Furuta and Hogan (1998) proposed a requirement for a factor(s) other than BMP4 secreted from the PR for the PLE to become lens.

**Fig. 9.**
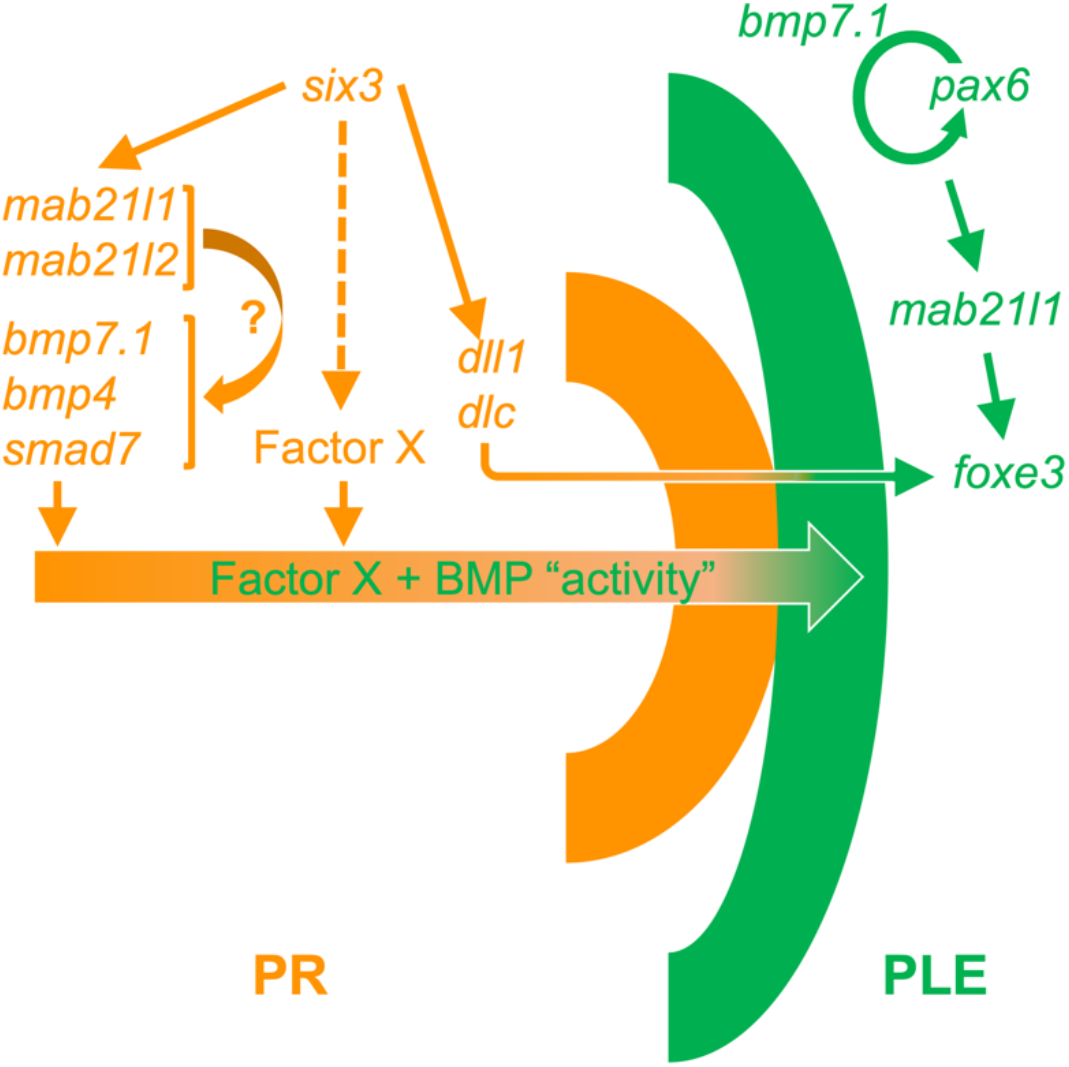
Model of cell non-autonomous Six3 function in the presumptive retina (PR) during neurulation. Genes shown in green are expressing in the PLE (thick green line) and genes in orange are expressing in the PR (thick orange line). Part of a gene hierarchy required for lens (arrows from *six3*) formation in the PLE known to date is summarized on right side, new interactions shown on left side of the figure. Dotted line shows potential role of unknown factors (Factor X) that also could be involved in mediating lens formation from the retina.

Another possible gene linking *six3* to BMP signaling in the retina is *lhx2* since previous studies have implicated the involvement of Lhx2 in regulating BMP signaling activity: *Lhx2* knockout mice showed that expression of *Bmp4* and *Bmp7* in the developing retina are downregulated, and that pSmad1/5/9 level is decreased in the ectoderm, suggesting that, as a result, the lens fails to form (Yun et al., 2009). Consistent with above possible linkage, we have observed that *lhx2* expression is reduced in the *six3* mutant retina at st.21 (data not shown). A further putative candidate for key signals in the optic vesicle is FGF, which is suggested to be essential for lens formation in a dose-dependent manner (Lovicu and McAvoy, 2005), though it is not known if FGF signaling is under the control of Six3. The possible connection of Six3 and Lhx2 and/or FGF signaling will need to be further investigated in the future.

As also noted in Fig. 9, and discussed earlier, *pax6* appears to have a primary influence on lens formation within the developing PLE (Fujiwara et al. 1994) where BMP signaling (via BMP7) is also required within the PLE (Wawersik et al., 1999). However, we also recognize that *six3* may have an autonomous effect in lens formation since transplants of WT PLE to mutant PR hosts did show positive crystallin expression in some embryos (Fig. 4). This possible role will need to be further investigated in the future.

We also found signaling networks are down regulated by reductions in nuclear proteins *mab21l1* and *mab21l2* in the *six3* mutant PR, both of which were shown as well to cause lens defects in knockout mice (Yamada et al., 2003; Yamada et al., 2004). Importantly, *Mab21l2* is known to cause a PR defect leading to lens defects (Yamada et al., 2004), which is consistent with our hypothesis that failure of the PR in frog *six3* mutant causes the lens defect seen here. By contrast, *Mab21l1* is suggested to function in the PLE cell-autonomously for lens formation; however, it is worth pointing out that the mutant mice also have abnormal retinal development (Yamada et al., 2003), possibly augmenting the failure of lens formation non-autonomously. Our data shows that *mab21l1* mRNA could partially rescue lens formation in the *six3* mutant as evidenced by *foxe3* and *cryg1* expression and coincident with recovery of BMP activity as evidenced by *smad7* expression, a readout of activation of BMP pathway signaling (Fig. 8). The mechanism of how *mab21l1* rescues BMP activity is currently unclear and needs to be investigated further.

Although we have previously shown that *foxe3* expression is under the control the Notch signaling pathway via *dll1* and *dlc* (Ogino et al., 2008), we did not observe rescue of *dll1* and *dlc* expression by *mab21l1* mRNA injection (data not shown), suggesting that the rescue of *foxe3* expression in the *six3* mutant by *mab21l1* mRNA (Fig. 8C) occurred independently from Notch signaling and thus *foxe3* appears to be regulated by multiple mechanisms. Consistent with this finding, the *foxe3* expression is still seen at st.21 but expression of *dll1* and *dlc* are broadly lost by st.21 in the *six3* mutant (Fig. 5).

As noted above, an important finding in our study is the independence of regulation of *pax6* and *six3* expression during early development, at least through neural tube closure, further highlighting the essential, independent role of *six3* in eye development. Although this result may appear to deviate from previous findings in the mouse (Goudreau et al., 2002; Liu et al., 2006) which suggest an interplay between these two genes, our studies are focused on earlier stages (at the neural plate and early neural tube stages) while the mouse studies largely focus on interactions between *Pax6* and *Six3* at later stages. The differences observed could also be due to species specific variations or technical challenges of conditional knockout systems involving Le-Cre, necessitated in the mouse in order to focus specifically on retina and lens formation (Dorà et al., 2014; Pathania et al., 2016; Song and Palmiter, 2018).

In summary, using the frog *six3* mutant which has eye defects but therefore has an advantage over the headless mouse *Six3* mutant for more direct study of eye development alterations, we have proposed novel evidence that Six3 functions cell non-autonomously in the developing PR to allow the overlying PLE to give rise to the lens through Mab21l1/Mab21l2 and BMP signaling. These processes are independent of *pax6* expression and are a key part of the mechanism determining lens fate occurring during neurula stages (Fig. 9).

## MATERIALS AND METHODS

### Frogs

The frogs used for this study were from the *X. tropicalis* colony maintained at the University of Virginia. All procedures involving frogs were performed in accordance with IACUC protocols approved by the University of Virginia. Embryos were staged according to Nieuwkoop and Faber, 1967. The wild type frogs used are from our in-house breeding stocks. These are not highly inbred but originated from the same ancestors as the Nigerian inbred line that was used for genome sequencing (Hellsten et al., 2010). The *six3* mutant frogs were offspring of the first CRISPR/Cas9 gene edited *six3* mutant F0 frogs (Nakayama et al., 2013). The *pax6* mutants were described before, and here we used a line having a 13-bp deletion in exon 7 (*pax6*^*ex7Δ13bp*^, Nakayama et al., 2015). The *six6* mutant (a HD domain mutation, *six6*^*Δ16bp complex*^, Fig. 1) was made using the target site as described below and in the Results section.

### Generation of mutant lines

The procedure of CRISPR/Cas9 gene editing in terms of a strategy of design of target sites, making PCR-based sgRNA templates, and synthesizing sgRNAs has been described previously (Blitz and Nakayama, 2022; Nakayama et al., 2013; Nakayama et al., 2014). In the early stage of this project, Cas9 mRNAs (Nakayama et al., 2013; Nakayama et al., 2014) were used, but later studies utilized Cas9 protein (CP01, PNA BIO INC). When Cas9 protein was used we incubated the Cas9 protein/sgRNA mixture at 37°C for 5 minutes before injection. Microinjection was done as described previously (Nakayama et al., 2013; Ogino et al., 2006). The *six6* target used in this study is as follows (shown as sense-strand sequence): GGGAACTCGCCCAAGCGAC(TGG) where the PAM sequence is in parentheses.

### Genotyping of embryos

Genotyping of *six3*^*Δ19bp*^, *six6*^*Δ16bp complex*^, and *pax6*^*ex7Δ13bp*^, a sequencing-free genomic PCR-based genotyping (in short SFG for sequencing-free genotyping) was done using the embryo lysate prepared as described before (Blitz and Nakayama, 2022; Nakayama et al., 2013; Nakayama et al., 2014), the concept for which is summarized in Fig. S3 using the *six3*^*Δ19bp*^ locus as an example. The *six3* genotyping for *six3*^*Δ12bp*^ and *six3*^*Δ6bp*^ was done as originally described followed by sequencing (Nakayama et al., 2013) since SFG has not yet been established for those lines. Primers for genotyping are listed in Table S1.

### *Whole–mount in situ* hybridization

Whole-mount *in situ* hybridization (WISH) with a modification for genotyping was carried out as described (Fish et al., 2014). All antisense probes were labeled with digoxigenin (Roche) and detected by BCIP/NBT or BM-Purple (Roche). The probes described previously are as follows: *six3, pax6, mab21l12* (Fish et al., 2014); *mab21l1* (Nakayama et al., 2015); *nrl, mafb* (Jin et al., 2012); *smad7* (Nakayama et al., 1998); *cryg1* (Offield et al., 2000). Newly described probes made from cDNAs are *dll1* (purchased from Open Biosystems, clone ID: 7605672), *notch2* (purchased from Open Biosystems, clone ID: 7678933), and *cryba1* (purchased from Open Biosystems, clone ID: 9018035). Other new probes made from PCR-based templates (Fish et al., 2014) are *foxe3, dlc, bmp7*.*1, bmp4* as described in Table S2.

### mRNA injection

The *mab21l1* mRNA (using the same plasmid used for making WISH probe above) was made by standard methods (e.g., see Nakayama et al., 2013, Fish et al., 2014). As illustrated in Fig.8A, *mab21l1* mRNA (500 pg per blastomere) was injected together with lineage tracer into two dorsal (animal) blastomeres at 4(8)-cell stage(s). We used only properly injected embryos, i.e., lineage tracer is seen in both left (white L) and right (white R) anterior region (Fig. 8A), but not only in one side (e.g., whole right half side, red R) which could occur due to inaccurate identification of dorsal blastomeres.

### Tissue transplantation

Embryos obtained by crossing *six3* heterozygous frogs were injected with 60 ng fluorescent dextran (Invitrogen, D1845) per embryo to provide labeled donor tissues (see Fig. 4A). As for host embryos, we used sibling embryos or independent WT embryos as needed. All embryos were raised at 22°C to st.15 in 0.1 x MBS, pH 7.5 (Nakayama and Grainger, 2022). Same stage host and donor embryos were selected, and their vitelline membranes removed with fine forceps. The embryos were lined up in host-donor pairs in adjacent wells in a clay-lined dish containing 1x MBS + 0.01% BSA a few at a time, and PLE’s were removed using tungsten needles and discarded from the host embryo, and then its donor partner’s PLE was removed and inserted into the host embryo. Transplants were held in place by small clay “fingers.” The transplants were allowed to heal for at least 15 minutes before host and donor embryo pairs were transferred from the operating dish into adjacent wells of a 24-well culture dish containing 0.5x MBS + 0.01% BSA. All operations were done on the embryos’ left sides, leaving the right sides as unoperated controls (see Fisher and Grainger, 2019 for further details). Transplant host and donor embryos were returned to the 22°C incubator and raised to st.37/38. At this point all embryos were fixed for 90 minutes in MEMFA [100 mM MOPS (pH 7.4), 2 mM EGTA, 1 mM MgSO_4_, 3.7% (v/v) formaldehyde], washed in PBS and donor embryos and tails from host embryos were digested for SFG (see above), while the heads of the hosts were dehydrated to 100% EtOH and stored in individual wells of a 96-well dish at -20°C until genotyping was complete and sufficient numbers of embryos were accumulated to analyze by WISH for *cryg1* expression.

### Immunostaining for phosphorylation of Smad1/9

Embryos were fixed at st.24 for 1 hour in 4% Paraformaldehyde in PBS then dehydrated to 100% MeOH and stored at -20°C. Embryos were then cut into anterior and posterior halves. The posterior halves were digested for SFG to identify mutant and wild type individuals as described above. The anterior half of each embryo was held in 100% EtOH at -20°C until genotyping was completed at which time, embryo heads of like genotype were pooled and embedded in paraffin (Paraplast Plus). Sections (10μm) were mounted onto gelatin-subbed slides and dried on a warming tray overnight (37°C).

Sections were dewaxed with xylene and rehydrated through graded ethanol washes into PBS and then treated with Vector Laboratories Antigen Unmasking Solution (VUS, citrate-based). A working solution was prepared from the 100x stock provided and then preheated (75ml in a large glass petri dish) in a microwave for 45 seconds on high, slides were then quickly added (4 slides at one time), the VUS reheated for 30 seconds at a low power level and allowed to sit undisturbed for an additional 60 seconds. Slides were quickly transferred to racks in cool (room temperature, RT) water for rinsing and then transferred back to PBS.

All slides were washed 2 x for 5 minutes in PBS/0.1%Tx-100 followed by blocking in 10% NGS/PBS-Tx for 1 hour at RT. Blocking solution was replaced with 1° antibody (Ab) (Phospho-Smad1/5/9 Rabbit mAb #13820, Cell Signaling Technology 13820T, 1:300) in 10%NGS/PBS-Tx and the slides were incubated overnight at 4°C. The following day slides were washed 5 x for 5 minutes in PBS-tx and then incubated in 2° Ab (GARGG-Alexafluor488, 1:300) overnight at 4°C. Finally, slides were washed 2 x for 5 minutes in PBS-tx and 3 x for 5 minutes in PBS and cover slipped with DAPI Fluoromount-G (SouthernBiotech).

### Image analysis of immunostained slides

Images obtained from Zeiss Axioskop equipped with a Zeiss AxioCam Mrc5 CCD camera were processed using Fiji v2.0.0-rc-69 (Schindelin et al., 2012). Ten manually selected regions of equal area generated by using the rectangular marker tool in Fiji. Regions from the PLE, PR, and the pharyngeal endoderm region of the foregut were selected, and Mean Gray Values were calculated for each of the 10 regions per section. The data pooled from four individual batches of immunostaining of each mutant and WT were analyzed using GraphPad Prizm v8.4.3 for Macintosh (GraphPad Software, San Diego, CA, USA) for comparison of Mean Gray Values of lens, retina, and the foregut between *six3* mutant and WT.

## Acknowledgements

We would like to acknowledge Jan Christian for *smad7* cDNA, and Ira Blitz for discussion about BMP signaling, Cristina D’Ancona for her assistance in CRISPR-targeting *six6* loci, and Merly Konathapally, Cameron Jackson, Sejal Kapoor, Sophie Tippit and Himabindu Gonugontla for assistance in imaging and performing WISH. We also thank Elizabeth Hensel for work genotyping animals and for input regarding statistical analysis.

## Competing interests

The authors declare no competing or financial interests.

## Author contributions

Conceptualization: S.M., T.N, R.M.G.; Methodology: S.M., T.N, M.F., R.M.G.; Formal analysis: S.M., T.N, M.F., R.M.G; Investigation: S.M., T.N, R.M.G.; Writing - original draft, review & editing: S.M., T.N., M.F., R.M.G.; Supervision: R.M.G.; Funding acquisition: R.M.G.

## Funding

This work was funded by NIH grant EY022954, research awards from the Sharon Stewart Aniridia Trust and Vision for Tomorrow, and research support from the Department of Ophthalmology, University of Virginia.

## SUPPLEMENTARY DATA

**Fig. S1.**
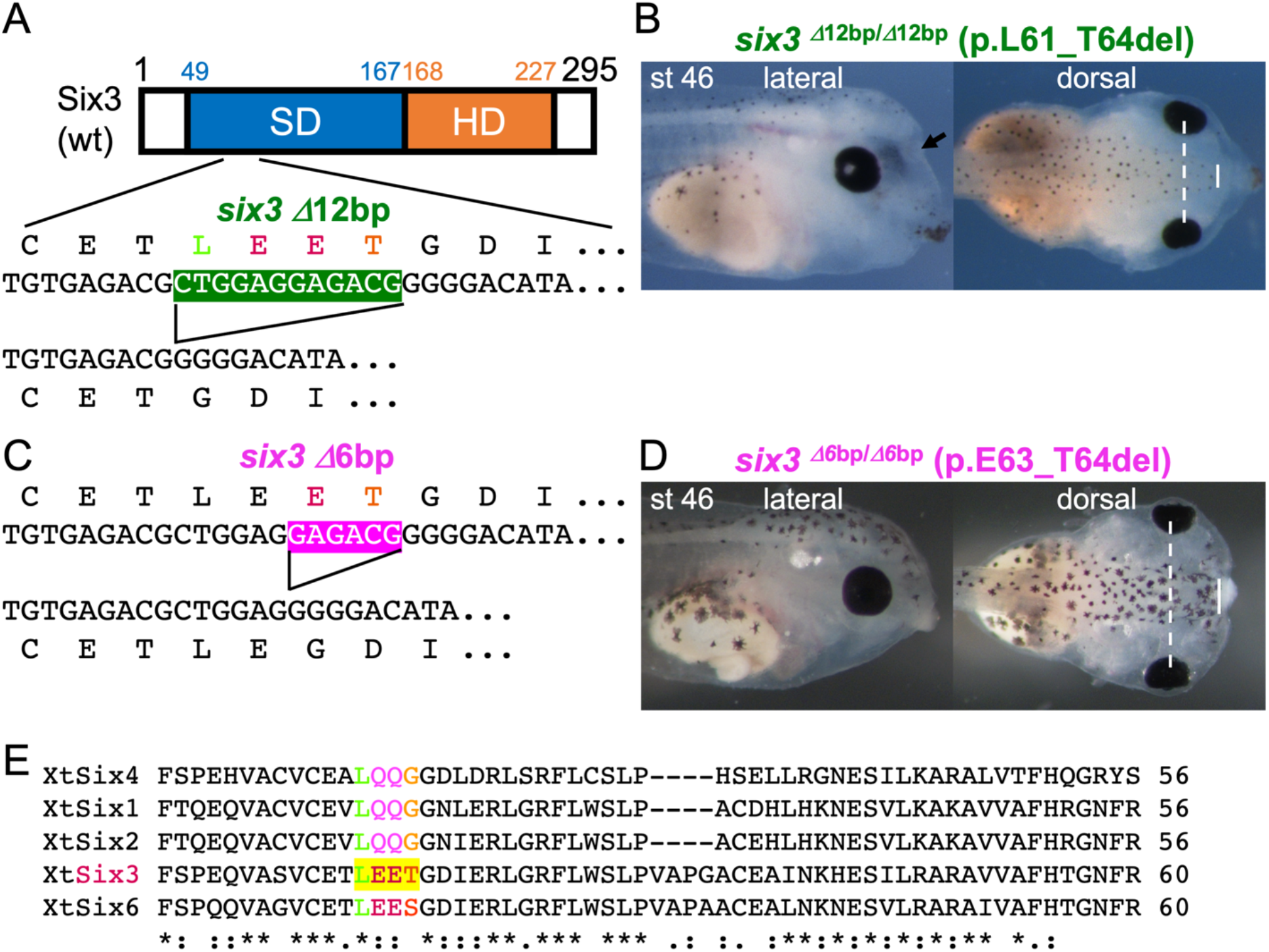
A hypomorphic mutation of *six3* gene in *X. tropicalis* causes a milder HPE and eye defects. (A) Structure of *six3* D12bp mutation. The resultant protein loses amino acids 61-64. (B) Phenotype of *six3*^D12bp/D12bp^ mutant, where the black arrow indicates a fused nasal pit. The eye phenotype is milder than null mutant (compare with Fig.1), but this mutation is still embryonic lethal. (C) Structure of *six3* D6bp mutation, losing amino acids 63-64. (D) Phenotype of *six3*^D6bp/D6bp^ mutant, which is indistinguishable from wild type at least during tadpole stages. White dotted lines indicate width between eyes and white solid lines width of the forebrain (B, D). (E) ClustalW alignment of parts of SDs of *X. tropicalis* SIX family members. Yellow-highlighted portion is lost in the *six3* D12bp and partially lost in the *six3* D6bp mutation, suggesting an important domain conserved in the SIX family.

**Fig. S2.**
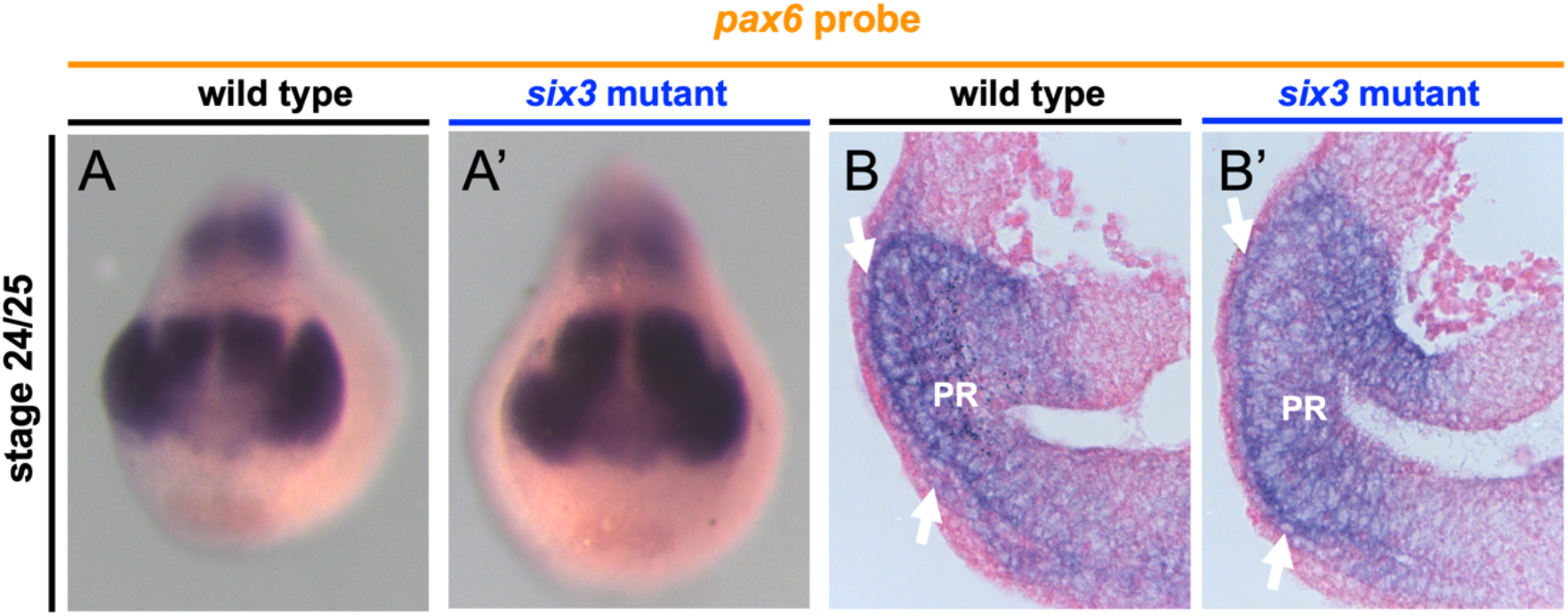
*pax6* expression in the eye is not affected in the *six3* mutant even at early tailbud stage. (A, A’) WISH images of wild type (A) and *six3* mutant (A’). (B, B’) Images of sectioned embryos of the same embryos shown in (A, A’), showing *pax6* stain is present in the presumptive lens ectoderm (PLE – white arrows) and presumptive retina (PR) in wild type (B) and *six3* mutant (B’).

**Fig. S3.**
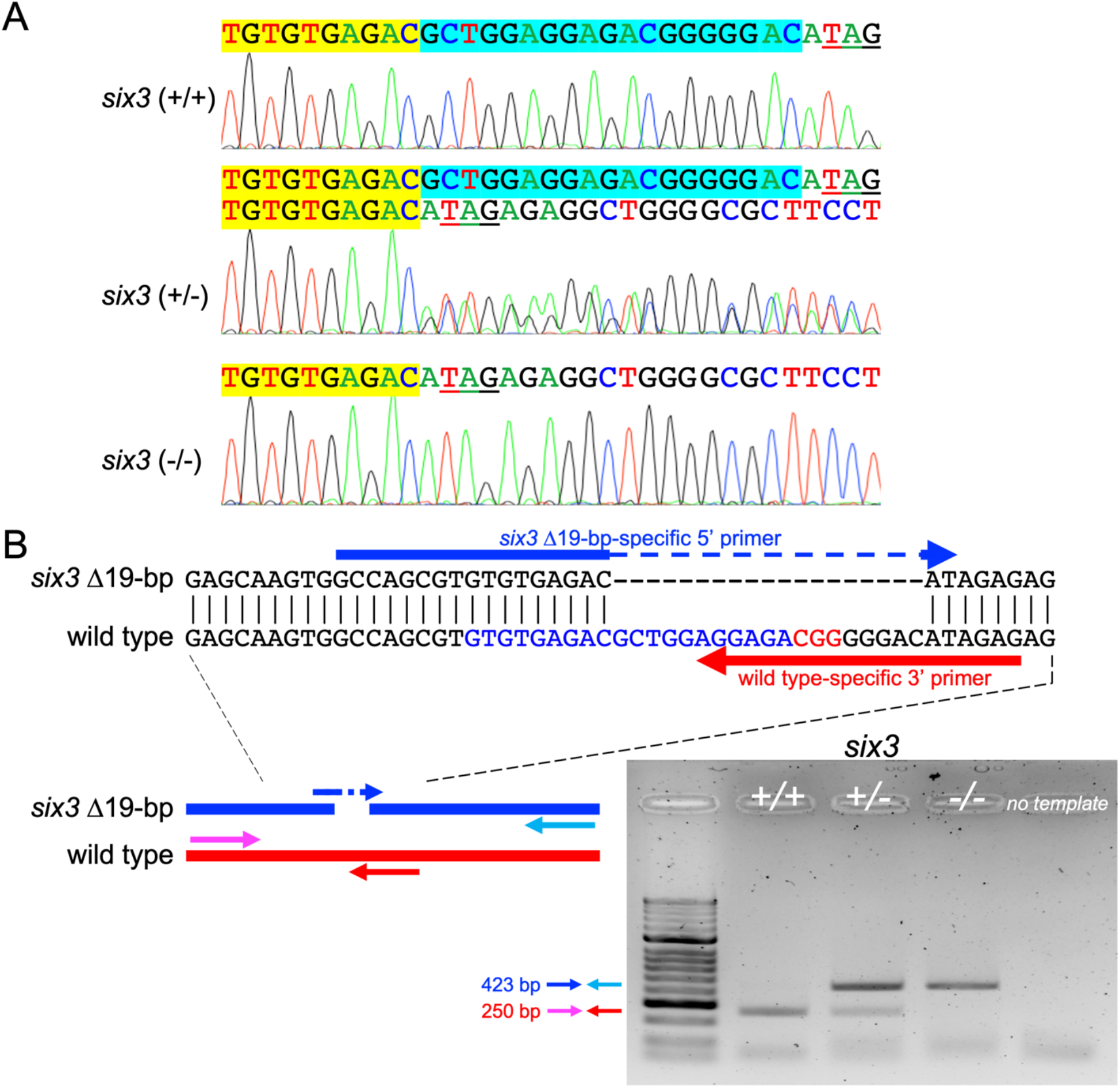
A concept of PCR-based genotyping. (A) DNA sequencing chromatograms of *six3* locus for wild type (+/+), heterozygote (+/-), and mutant (-/-) genotyped by the regular method. The yellow-or cyan-highlight is the same color code as used in Figure 1. (B) PCR-based genotyping. The top panel shows examples of designs of the PCR primers specific to mutation (blue) and wild type (red) *six3* loci, where gaps (shown by -) are inserted for alignment. The bottom left shows the schematic PCR design, where cyan or magenta primers are used respectively with blue (to amply the mutant locus) or red primers (to amply the wild type locus). The right panel shows the results of PCR for wild type (+/+), heterozygote (-/+), or mutant (-/-).

**Table S1.**
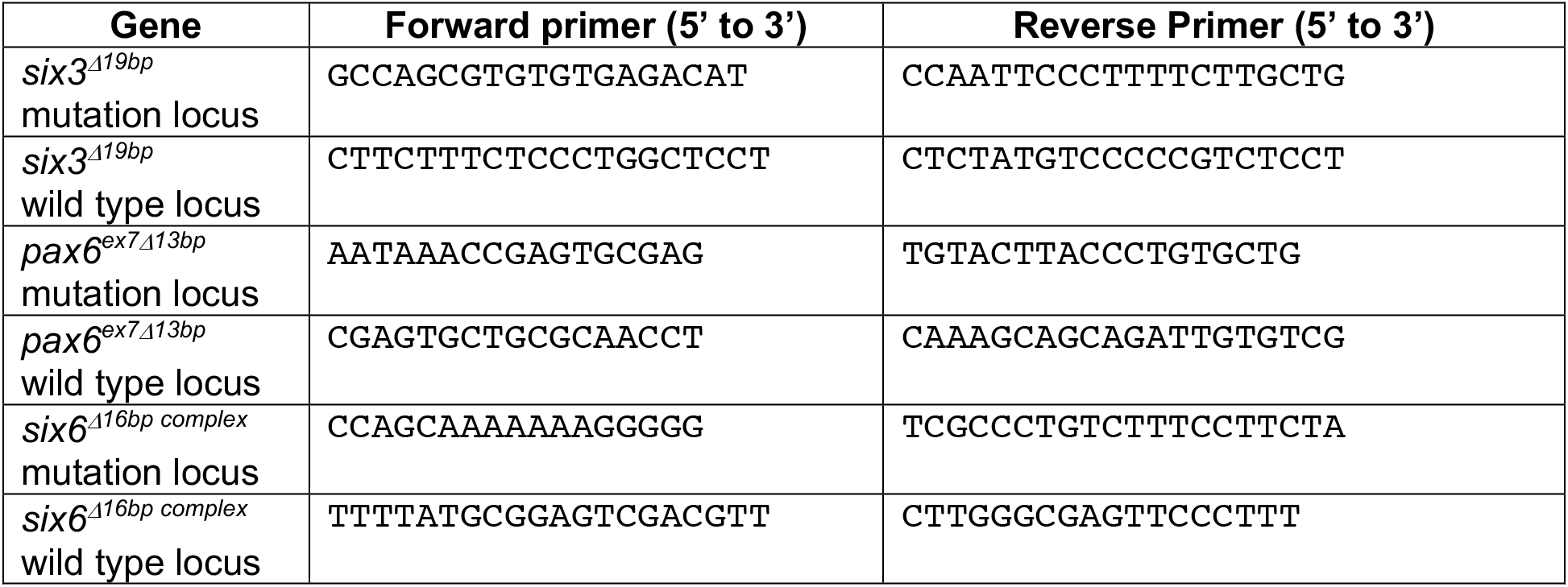
Primer list for genotyping.

**Table S2.**
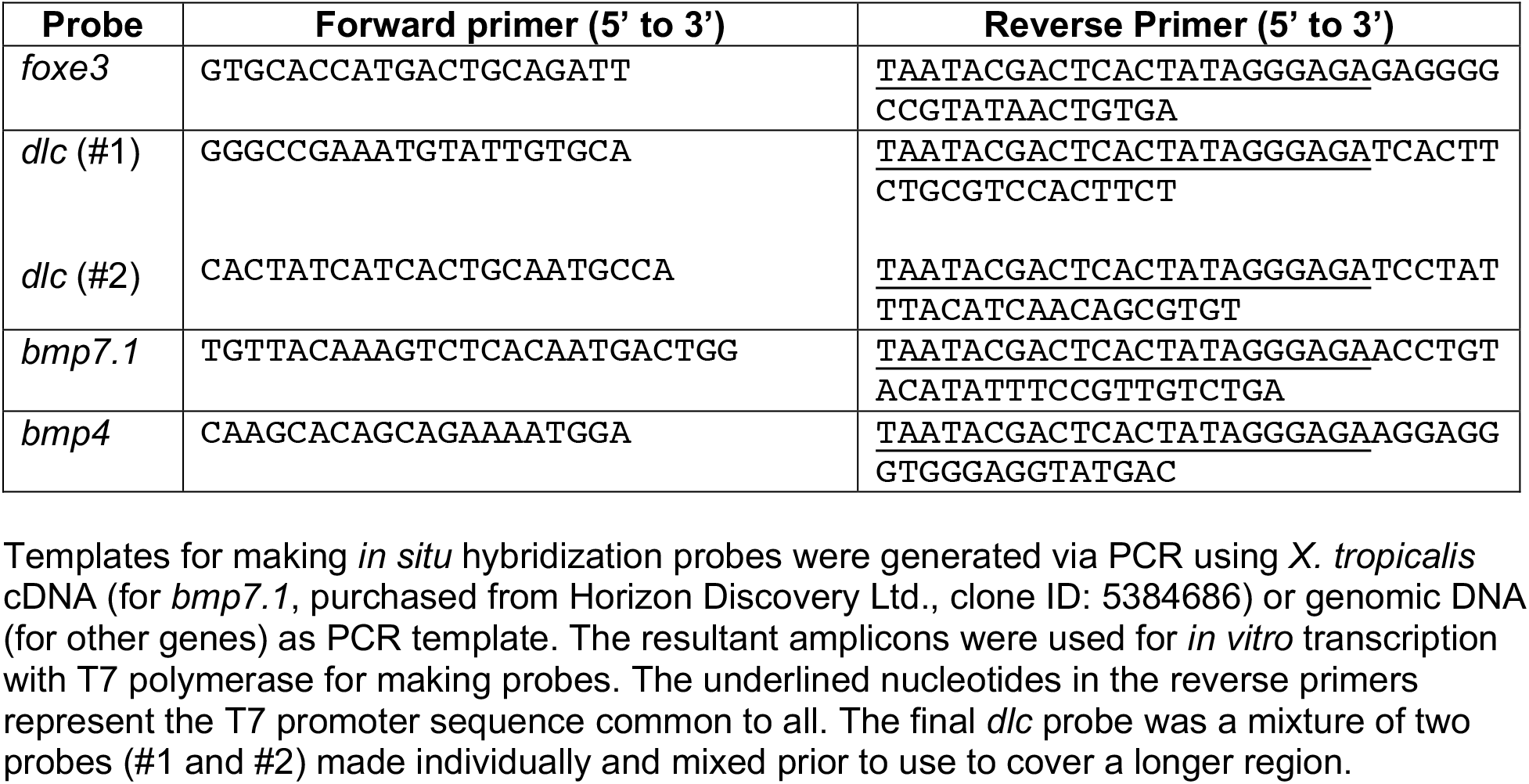
*in situ* hybridization probes made by using PCR products as templates.

## Notes

### Competing Interest Statement

The authors have declared no competing interest.

